# Distinct contributions of three GABAergic interneuron populations to a mouse model of Rett Syndrome

**DOI:** 10.1101/155382

**Authors:** James M. Mossner, Renata Batista-Brito, Rima Pant, Jessica A. Cardin

**Author notes:** Corresponding author: Jessica A. Cardin, 333 Cedar St., PO Box 208001, New Haven CT 06520.

## Abstract

**Background:** Rett Syndrome is a devastating neurodevelopmental disorder resulting from mutations in the gene MeCP2. MeCP2 is a transcriptional regulator active in many cell types throughout the brain. However, mutations of MeCP2 restricted to GABAergic cell types largely replicate the behavioral phenotypes associated with mouse models of Rett Syndrome, suggesting a key role for inhibitory interneurons in the pathophysiology underlying this disorder.

**Methods:** We generated conditional deletions of MeCP2 from each of three major classes of GABAergic interneurons, the parvalbumin (PV), somatostatin (SOM), and vasoactive intestinal peptide (VIP)-expressing cells, along with a pan-interneuron deletion from all three GABAergic populations. We examined seizure incidence, mortality, and performance on several key behavioral assays.

**Results:** We find that each interneuron class makes a contribution to the seizure phenotype associated with Rett Syndrome. PV, SOM, and VIP interneurons made partially overlapping contributions to deficits in motor behaviors. We find little evidence for elevated anxiety associated with any of the conditional deletions. However, MeCP2 deletion from VIP interneurons causes a unique deficit in marble burying. Furthermore, VIP interneurons make a distinct contribution to deficits in social behavior.

**Conclusions:** We find an unanticipated contribution of VIP interneuron dysfunction to the MeCP2 loss-of-function model of Rett Syndrome. Together, our findings suggest a complex interaction between GABAergic dysfunction and behavioral phenotypes in this neurodevelopmental disorder.

## Introduction

Rett Syndrome is a severe neurodevelopmental disorder caused by mutations in the gene encoding methyl-CpG binding protein 2 (MeCP2). Children affected by Rett develop normally during the initial year of postnatal life but rapidly regress thereafter, showing loss of motor skills and language and developing cognitive impairments, ataxia, respiratory problems, and stereotyped hand movements (1). Extensive previous work has shown that many Rett-associated phenotypes are replicated in MeCP2 loss-of-function mouse models (2-6). Rett Syndrome is strongly associated with seizure (1, 7, 8), suggesting a possible role for GABAergic dysregulation in the pathophysiology underlying these symptoms. Indeed, previous work in mice found that conditional mutations of MeCP2 restricted to GABAergic neurons recapitulate most of the observed phenotypes (2, 5) whereas rescue of MeCP2 solely in GABAergic neurons ameliorates many phenotypes (9), suggesting a key role for GABAergic dysregulation in Rett Syndrome.

One major challenge in exploring GABAergic dysfunction in Rett Syndrome is the diversity of inhibitory interneurons, which can be subdivided into distinct classes with different physiology, synaptic targets, and molecular markers. The functions of three major classes of GABAergic cells have recently been explored in depth: fast-spiking basket cells that co-express the calcium-binding protein parvalbumin (PV) and target the cell bodies of excitatory neurons, low-threshold spiking cells that co-express the peptide somatostatin (SOM) and target the distal dendrites of excitatory neurons, and sparse interneurons that co-express vasoactive intestinal peptide (VIP) and mainly target other inhibitory interneurons. These three classes are thought to play distinct roles in regulating local circuit activity throughout the brain (10) and may thus make different contributions to the pathophysiology underlying Rett Syndrome.

Recent work has suggested that MeCP2 loss of function mutations in PV and SOM interneurons may have distinct consequences. Conditional deletion of MeCP2 from PV interneurons caused motor, sensory, cognitive, and social deficits, whereas deletion from SOM interneurons resulted in increased behavioral stereotypy and late-onset seizures (5). However, Cre-dependent MeCP2 deletion in PV and SOM cells is not well matched developmentally, as the SOM promoter is active during early embryonic development but the PV promoter is not active until after the first two postnatal weeks (11). Furthermore, nothing is known about the relative contribution of VIP interneurons to this disorder.

Here we directly compared the impact of MeCP2 loss-of-function in PV, SOM, and VIP interneurons. We generated conditional mutations of MeCP2 in each interneuron class and compared these with a conditional pan-interneuron mutation that includes all three interneuron classes and drives embryonic deletion. To identify the distinct contributions of each interneuron class, we assayed seizure incidence and mortality, locomotor and anxiety phenotypes, and social behavior. We find that PV, SOM, and VIP interneuron deficits may each contribute to seizure incidence following MeCP2 mutation. All three interneuron classes likewise contribute to general locomotor deficits. However, MeCP2 deletion from VIP, but not PV or SOM, interneurons caused deficits revealed by marble burying. Furthermore, MeCP2 deletion from VIP interneurons made a unique contribution to social interaction impairments. Overall, our findings suggest an unanticipated role for VIP interneuron dysfunction in the MeCP2 model of Rett Syndrome. However, partially overlapping contributions from the three major interneuron classes suggest that these GABAergic populations may play intersecting roles in neurodevelopmental pathophysiology.

## Material and methods

### Animals

All experiments were approved by the Institutional Animal Care and Use Committee of Yale University. We used the Dlx5/6-Cre (JAX#008199; (12)), PV-Cre (JAX#008069; (13)), SOM-Cre (JAX#013044; (11)), and VIP-Cre (JAX#010908; (11)) mouse lines to target all forebrain GABAergic interneurons, Parvalbumin-expressing interneurons, Somatostatin-expressing interneurons, and Vasoactive Intestinal Peptide-expressing interneurons, respectively. We crossed each Cre line to the conditional MeCP2 line (MeCP2^f/f^; JAX# 007177; (4)). In each case, we assayed male mice that were hemizygous for the floxed MeCP2 allele and heterozygous for Cre. In a subset of experiments, we compared the interneuron-specific crosses with male mice from the MeCP2 knockout line (MeCP2^−/y^; JAX#003890; (4)). All crosses were made on a C57BL/6J background (JAX#000664). All behavioral assays were performed at 4 months except for the Dlx5/6-cre^+/-^ MeCP2^f/y^ animals, which were assayed at 3 months due to early morbidity.

### Immunohistochemistry

For immunofluorescent staining of brain tissue, mice were perfused with 4% Paraformaldehyde and post fixed for an hour before transferring into successive sucrose solutions at 15 and 30%. 20μm thick cryosections were prepared for IHC. Tissue was incubated with 1.5% normal goat serum (NGS) (Life Technologies) and 0.1% Triton X-100 (Sigma) in PBS for 60 min at room temperature. Sections were incubated with primary antibodies [Rat Anti-Somatostatin 1:250 (Millipore MAB354); Anti-parvalbumin 1:1000 (Sigma P3088); Anti-VIP 1:250 (ImmunoStar 20077); Anti-MeCP2 1:250 (Millipore 07-013)] in the blocking buffer overnight at 4 °C. After washing three times with buffer, sections were incubated with secondary antibodies for 1 h at room temperature [Secondary antibodies: Alexa Fluor 488, 594 or 647 (Life Technologies, 1:1000)]. Finally, coverslips were mounted using ProLong Gold Mounting Medium with DAPI (Life Technologies) and imaged at 10x. Quantifications were performed in Adobe Photoshop. Pictures were divided into a grid measuring 1X1 cm in total and cells were counted in each grid square. The number of cells positive for antibody staining against MeCP2 was counted to assay the proportion of co-expressing cells.

### Extracellular recordings

LFP recordings were made with tetrodes (Thomas Recording GMBH, Germany) targeted to layers 2/3 and 5 of parietal cortex above the CA1 field of the dorsal hippocampus and from within the CA1 (AP: +1.5-2mm; ML: 1.2-1.75, Franklin and Paxinos, 2001). Signals were digitized and recorded with a DigitalLynx 4SX system (Neuralynx, Bozeman MT). All data were sampled at 40kHz and recordings were referenced to the cortical surface. LFP data were recorded with a bandpass 0.1-9000Hz filter.

### Seizure detection

All mice were handled for at least 10 min each day throughout the study. During the daily handling regime, the mice were assessed for seizures. If a seizure did occur, the mouse was immediately returned to its home cage. Seizures were defined as events reaching Racine scale levels 4 or 5, with animals exhibiting rearing and forelimb clonus or rearing, forelimb clonus, and falling.

### Morbidity analysis

After weaning at P21, all mice were monitored every day throughout the study. All deaths were noted, and animals were tracked daily until P500.

### Behavioral analysis

The open field, elevated plus maze, marble-burying, and sociability assays were performed under low-level (20-25 lux) red illumination. In each assay, mice were given 15 minutes to acclimate to the behavioral assay room. In all cases, the researcher was blind to the genotypes of the mice until after all behavioral data were scored.

#### Open field

The open field assay was performed in a 30cm square box divided into nine quadrants. Custom software (Labview) was used to control a camera recording the mouse’s path in the box. At the start of the session, the mouse was placed in the center quadrant of the box and allowed move freely for twenty minutes. After the time elapsed, the mouse was returned to the home cage and the box was cleaned for the next mouse. ImageJ software was used to analyze the total distance traversed by the mouse and the number of times the mouse crossed the center quadrant during the twenty-minute period. The mouse was deemed to have entered the center quadrant when all four feet were inside the quadrant and to have left the quadrant when all four feet were outside.

#### Elevated Plus Maze

Custom Labview software was used to control a camera recording the mouse’s locomotion on the maze. At the beginning of the session, the mouse was placed in the center of the maze and allowed to freely move on either arm for five minutes. At the end of the session, the mouse was returned to the home cage and the maze was cleaned for the next mouse. Video recordings of mouse behavior were hand-scored to determine the amount of time spent in the open and closed arms of the maze.

#### Marble Burying

12 marbles were evenly placed in a cage with 1 inch of clean bedding. Custom Labview software was used to control a camera recording the mouse’s activity in the cage. The mouse was placed in the center of the cage with the marbles and allowed to explore the cage for 20 minutes. At the end of the session, the mouse was returned to the home cage and the marbles were cleaned with a 10% bleach solution.

The proportion of marbles buried was analyzed in ImageJ using the “analyze particles” function to compare the initial and final exposed surface area of the marbles.

#### Sociability

The sociability apparatus was divided into three equal areas with Plexiglas dividers, each with an opening allowing access to neighboring chambers. Custom Labview software was used to control a camera recording the mouse’s activity in the chamber. An unfamiliar age-and sex-matched conspecific mouse (reared in separate cages, C57BL/6 genotype) was placed into a small cylindrical holding cage in one side of the chamber and an identical empty holding cage was placed in the other side. The location of the conspecific was randomly varied across trials. At the beginning of the session, the test mouse was placed in the central chamber and habituated to the central chamber for ten minutes. The dividers were then removed to allow the mouse to freely move among all the chambers for ten additional minutes. At the end of the session, the mice were returned to their home cages and the apparatus was cleaned. Video recordings of the mouse’s behavior were hand-scored to determine the amount of time spent in each of the three partitions and the number of approaches the test mouse made to the conspecific and the empty holding cage. An approach was defined as the test mouse coming within a 5-centimeter radius of a cage or making contact with a cage.

### Statistical analysis

Paired and unpaired non-parametric tests generated in GraphPad Prism (version 7 for Mac; San Diego CA) were used throughout the study due to non-normal data distributions. Exact p values are reported for all tests.

## Results

### MeCP2 expression in PV, SOM, and VIP interneurons

To confirm that MeCP2 is expressed in the three major populations of GABAergic interneurons, we co-stained sections of primary visual cortex from adult mice with antibodies for interneuron markers and MeCP2 (Fig. 1). As reported previously (5), nearly all PV and SOM interneurons expressed MeCP2. In addition, ~80% of VIP interneurons expressed MeCP2, suggesting a previously unappreciated role for this signaling pathway in VIP interneuron function.

**Figure 1.**
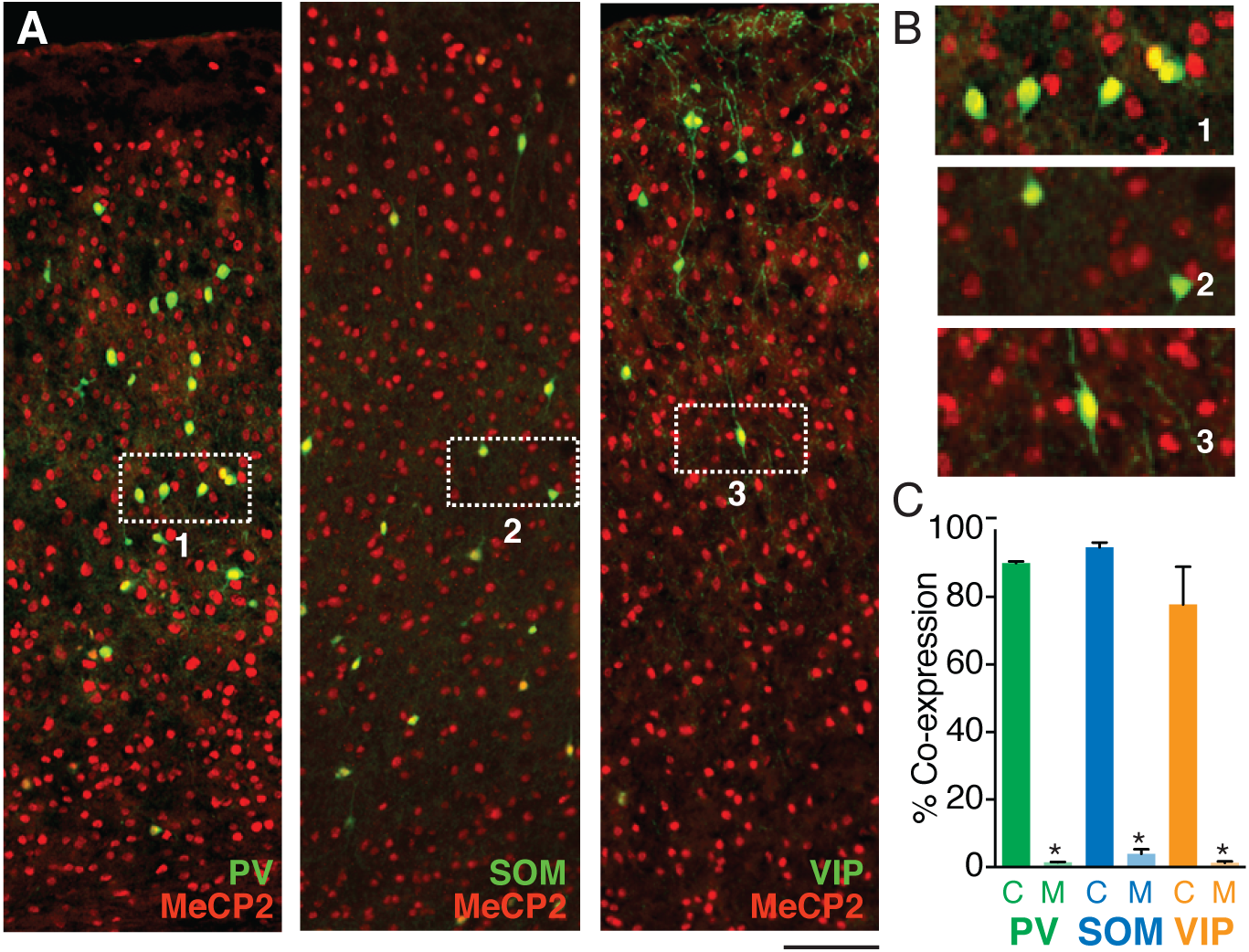
MeCP2 is expressed in three major GABAergic interneuron classes. A. Co-staining for interneuron markers (green) and MeCP2 (red) reveals a high degree of co-expression in PV (left), SOM (middle), and VIP (right) interneurons in the cortex. B. Expanded view of insets 1-3 from A. C. Crossing interneuron-specific Cre lines with the conditional MeCP2 line results in near-complete removal of MeCP2 expression from each target population. *p < 0.05

To determine the relative contributions of each interneuron class to behavioral phenotypes observed following MeCP2 deletion from all interneurons (2), we generated four lines of conditional deletion mice lacking MeCP2 specifically in PV (PV-Cre^+/-^ MeCP2^fly^; PV mutants), SOM (SOM-Cre^+/-^MeCP2^f/y^; SOM mutants), or VIP (VIP-Cre^+/-^ MeCP2^f/y^; VIP mutants) interneurons or in all three populations of GABAergic interneurons (Dlx5/6-Cre^+/-^MeCP2^fly^; Dlx5/6 mutants) (12, 14-16) by crossing MeCP2^f/f^ animals (17) to interneuron-specific Cre lines. We examined the efficacy of conditional removal of MeCP2 from targeted interneuron populations in the cortex of Cre^+^MeCP2^f/y^ animals (Fig. 1). Whereas 90.0 ± 0.6 % of SOM^+^ cells expressed MeCP2 in control animals, only 1.4 ± 0.1 % of SOM cells co-expressed MeCP2 in the SOM-Cre^+/-^MeCP2^f/^ animals (p = 0.029; n = 4 controls and 4 mutants; Mann-Whitney U test). Likewise, 94.7 ± 1.5% of PV^+^ cells in controls, but only 4.0 ± 1.4% in the PV-Cre^+/-^MeCP2^fly^ animals, co-expressed MeCP2 (p = 0.036; n = 4 controls and 4 mutants). Similarly, 77.8 ± 6.4% of VIP^+^ cells in controls and 1.2 ± 0.2% in the VIP-Cre^+/-^MeCP2^f/y^ animals co-expressed MeCP2 (p = 0.041; n = 4 controls and 4 mutants). These results suggest near-complete removal of MeCP2 expression in each of the targeted interneuron classes.

### Seizure incidence following MeCP2 deletion

Previous work identified a characteristic seizure phenotype resulting from MeCP2 deletion in the brain (2) and found that MeCP2 deletion from SOM-expressing cells may confer a late-onset tendency to seizure (5). We therefore evaluated the incidence of seizure in each of the three interneuron-specific MeCP2 deletion lines from weaning (P21) through late adulthood (P500). We compared the impact of MeCP2 deletion from PV, SOM, or VIP interneuron populations with simultaneous deletion from all three interneuron classes using the Dlx5/6-Cre line. We further compared the interneuron deletion mice with MeCP2^f/y^ littermate controls. To identify the relative impact of MeCP2 deletion from GABAergic cells compared to loss of MeCP2 in all cells, we also compared seizure incidence in the interneuron-specific deletion mice with that in male mice carrying a complete knockout of the MeCP2 gene (MeCP2^−/y^).

MeCP2 deletion from specific interneuron populations had markedly different effects on seizure incidence (Fig. 2A). We found that 100% of MeCP2^−/y^ (n = 10) and Dlx5/6 mutant (n = 13) mice exhibited at least one seizure, compared to only 52.9% of SOM mutants (n = 36), 35.0% of PV mutants (n = 22), and 37.5% of VIP mutants (n = 8). In comparison, the seizure rate in MeCP2^f/y^ animals was 17.1% (n = 38). The mean age of initial seizure was significantly earlier in MeCP2^−/y^ (41.3 ± 3.7 days; p = 0.009) and Dlx5/6 mutants (95.9 ± 10.1 days; p = 0.033), but not PV (108.9 ± 27.2 days; p = 0.464), SOM (149.4 ± 14.3 days; p = 0.99), or VIP (128.3 ± 38.0 days; p = 0.99) mutants, compared to MeCP2^f/y^ controls (162.3 ± 14.1 days; One-way Kruskal-Wallis test with Dunn’s multiple comparisons test; Fig. 2A-B). In a subset of SOM mutants, we performed electrophysiological recordings in the cortex and hippocampus of awake animals and confirmed spontaneous seizure activity in each animal at ~P200 (n = 4 mutants; Fig. 2C). These results suggest that MeCP2 deletions from PV, SOM, and VIP populations each contribute a degree of seizure pathophysiology.

**Figure 2.**
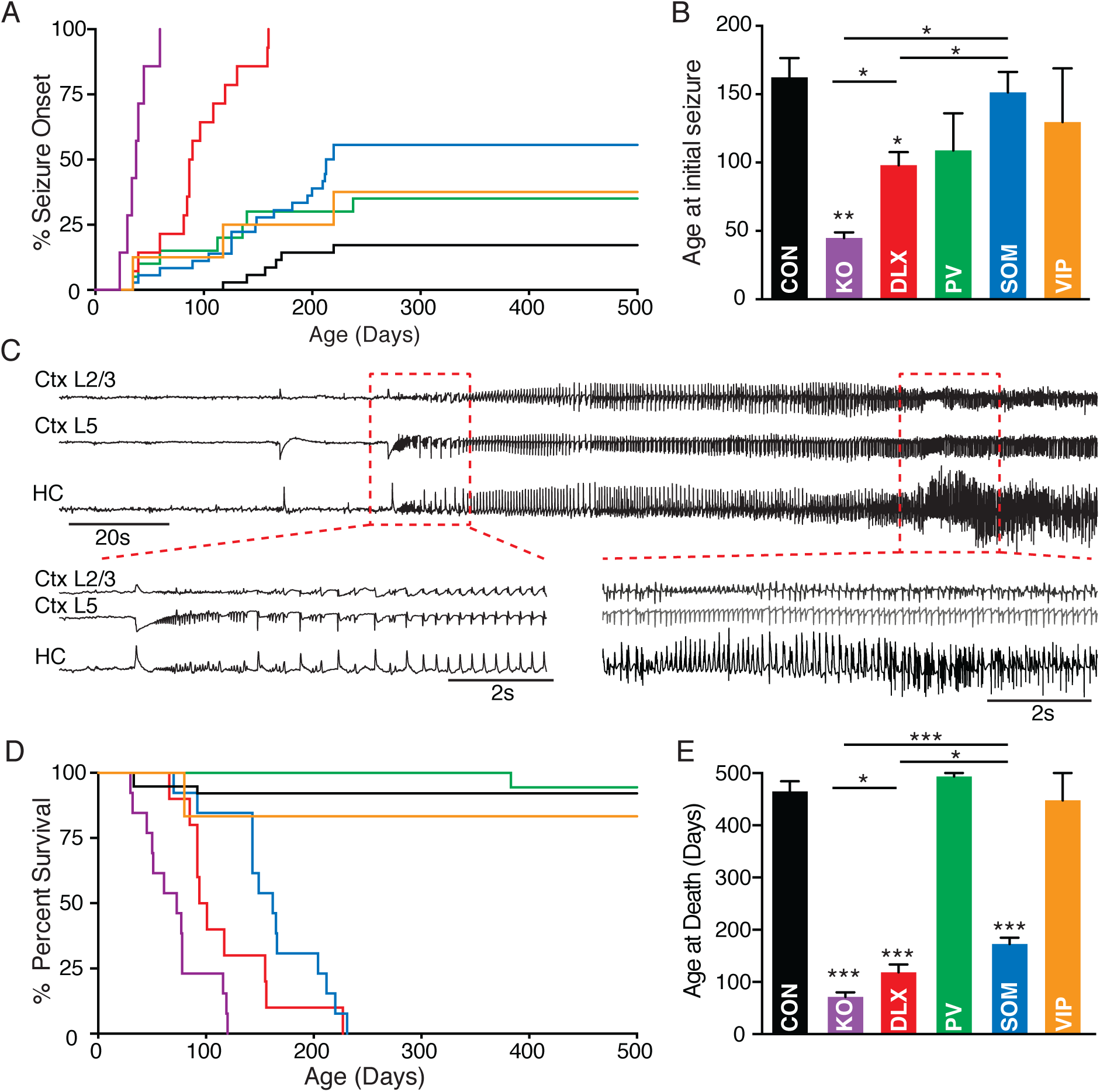
Seizure incidence following conditional deletion of MeCP2 from GABAergic interneurons. **A**. Cumulative distribution plot of the ages at which animals exhibited initial seizure activity for MeCP2^f/y^ controls (black; n = 38), MeCP2^−/-^(magenta; n = 10), and Dlx5/6 (red; n = 13), PV (green; n = 22), SOM (cyan; n = 36), and VIP (orange; n = 8) mutants. **B**. Mean age at initial seizure for each group shown in A. **C**. Example local field potential recordings from parietal cortical layers 2/3 and 5 and the CA1 of the hippocampus showing spontaneous ictal activity in an awake SOM-Cre^+/-^MeCP2^f/y^ animal at P220. **D**. Cumulative distribution plots of survival for controls (n = 38) and MeCP2^−/y^ (n = 13), Dlx5/6 (n = 10), PV (n = 18), SOM (n = 16), and VIP (n = 8) mutants. **E**. Mean age at death for each group shown in D. Death age was set to P500 for animals surviving beyond day 500. *p < 0.05, ***p < 0.0001

In addition to assaying the onset of seizure activity, we compared the incidence of death associated with seizure in each mouse line (Fig. 2D-E). We found that the mean age of death in MeCP2^−/y^ (71.5 ± 8.7 days; n = 13; p < 0.00001), Dlx5/6 mutants (118.5± 15.2 days; n = 10; p< 0.0001), and SOM mutants (172.5 ± 12.2 days; n = 16; p < 0.00001), but not PV (493.5 ± 6.5 days; n = 18; p = 0.88) or VIP mutants (447.5 ± 52.5 days; n = 8; p = 0.99), was significantly decreased compared to MeCP2^f/y^ controls (464.7 ± 19.9 days; n = 38; One-way Kruskal-Wallis test with Dunn’s multiple comparisons test; Fig. 2E). The mean age of death in SOM mutants was significantly elevated in comparison to either Dlx5/6 (p = 0.011) or MeCP2^−/y^ (p = 0.024; One-way Kruskal-Wallis test with Dunn’s multiple comparisons test) mutants. In addition, mean age of death was significantly later for Dlx5/6 mutants than for MeCP2^−/y^ animals (p = 0.038; One-way Kruskal-Wallis test with Dunn’s multiple comparisons test; Fig. 2E), suggesting that GABAergic interneuron dysfunction does not entirely account for the early seizure phenotype in the MeCP2 deletion model. These results suggest that although MeCP2 deletion specifically from PV, SOM, and VIP interneurons is associated with elevated seizure incidence in mature animals, only seizure activity following SOM-specific mutations is accompanied by increased mortality rates.

### Behavioral phenotypes associated with interneuron-specific MeCP2 deletion

We next characterized the contributions of MeCP2 deletions in each interneuron population to behavioral deficits. We found that locomotor activity in the open field test was impaired in pan-interneuron Dlx5/6 mutants (4988 ± 648.8 cm; n = 7; p = 0.011) and in the PV- (5493 ± 461.7 cm; n = 8; p = 0.021) and VIP- (5216 ± 563.7 cm; n = 8; p = 0.043) specific lines, but not the SOM mutants (6136 ± 575.8 cm; n = 13; p = 0.06), compared to controls (7912 ± 567.2; n = 18; Kruskal-Wallis multiple comparisons test; Fig. 3A). Center crossings in the open field, a measure of anxiety (18), were also impaired in the Dlx5/6 (19.7 ± 5.4 crosses; n = 7; p = 0.002), PV (26.0 ± 3.7 crosses; n = 8; p 0.033), SOM (25.1 ± 2.9 crosses; n = 13; p = 0.001), and VIP (16.6 ± 4.5 crosses; n = 8; p = 0.002) mutants as compared to controls (40.8 ± 2.0 crosses; n = 18; KruskalWallis multiple comparisons test; Fig. 3B). In contrast, we found no significant impairments in percentage of time spent in the open arms of the elevated plus maze task, another measure of anxiety, for the Dlx5/6 (39.0 ± 5.6%; n = 7; p = 0.99), PV (39.7 ± 6.3%; n = 8; p = 0.99), SOM (34.0 ± 3.4%; n = 13; p = 0.99), or VIP (33.8 ± 5.7%; n = 8; p = 0.99) mutants compared to controls (37.8 ± 5.4%; n = 18; Kruskal-Wallis multiple comparisons test; Fig. 4A). These results suggest that locomotor deficits are a common outcome of GABAergic dysregulation following MeCP2 deletion. We further found a significant impairment in marble burying, a measure of anxiety and exploratory motor function, in the pan-interneuron Dlx5/6 mutants (7.0 ± 4.6 marbles; n = 7; p = 0.014) compared to the controls (30.5 ± 3.6 marbles; n = 16; Kruskal-Wallis multiple comparisons test). This deficit was fully replicated by MeCP2 deletion specifically from VIP (10.9 ± 0.9 marbles; n = 8; p = 0.016), but not PV (26.3 ± 4.9 marbles; n = 7; p = 0.99) or SOM (20.1 ± 2.5 marbles; n = 13; p = 0.083), interneurons (Fig. 4B). VIP interneurons may thus make a cell type-specific contribution to the deficits in marble burying observed following MeCP2 deletion from all GABAergic neurons (2).

**Figure 3.**
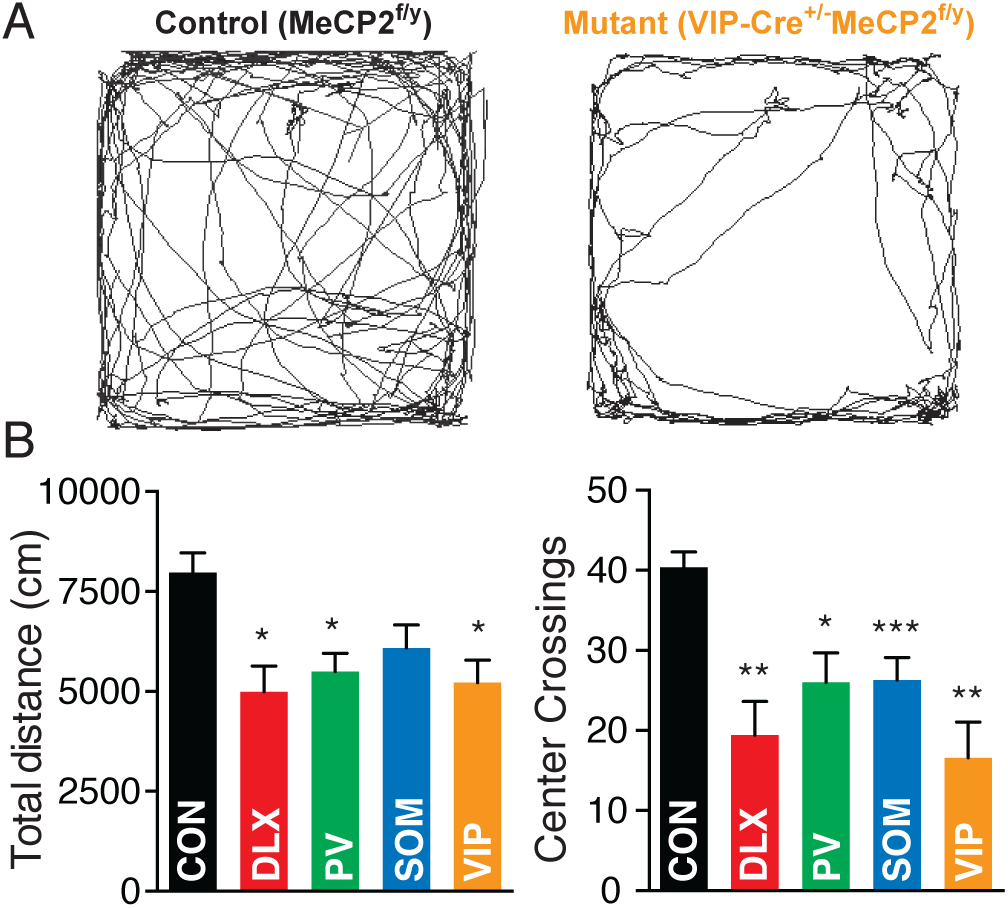
MeCP2 deletion disrupts locomotor behavior. **A**. Example open field trajectories for an MeCP2^f/y^ control (left) and a VIP mutant (right). **B**. Mean distance traveled (left) and number of center crossings (right) for controls (n = 14) and Dlx5/6 (n = 7), PV (n = 8), SOM (n = 13), and VIP (n = 8) mutants. *p < 0.05, **p < 0.01, ***p <0.001

**Figure 4.**
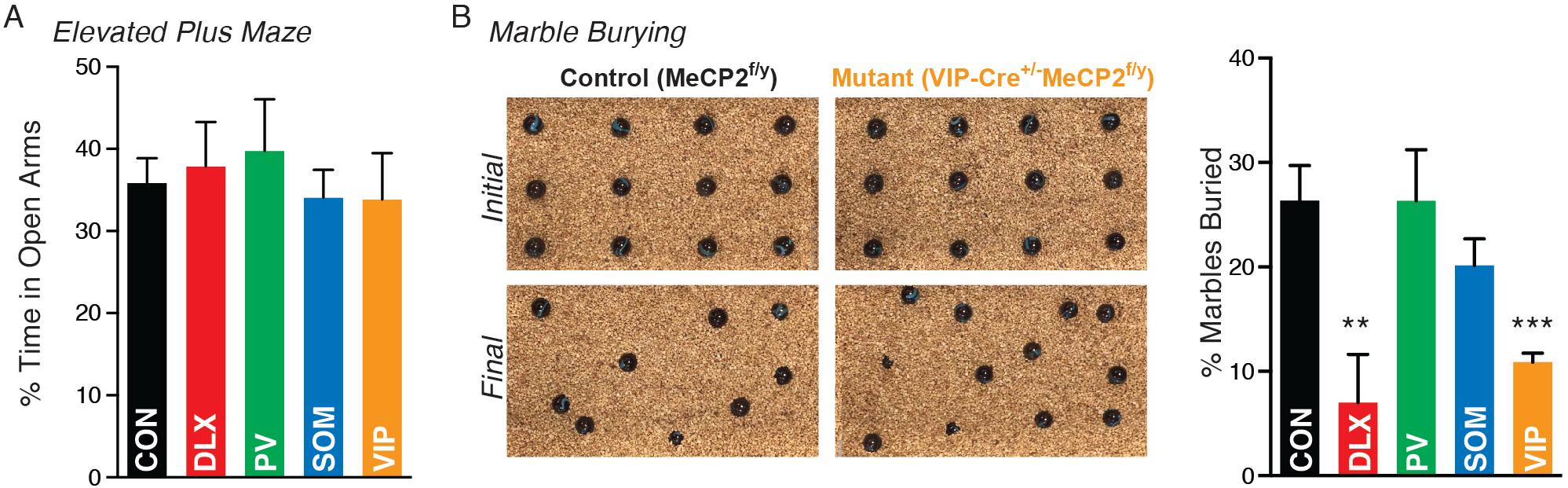
Loss of MeCP2 in VIP interneurons contributes to disrupted marble burying. A. Mean time spent in the open arms of the elevated plus maze for MeCP2^f/y^ controls (n = 14) and Dlx5/6 (n = 7), PV (n = 7), SOM (n = 13), and VIP (n = 6) mutants. B. Left: Example pictures of marbles buried by a control (left) and a VIP mutant (right). Right: Mean percentage of marbles buried by controls (n = 14) and Dlx5/6 (n = 7), PV (n = 7), SOM (n = 13), and VIP (n = 8) mutants. *p < 0.05, **p < 0.01, ***p <0.001

Because impaired social interactions are a hallmark of autism spectrum disorders and have been previously observed in the MeCP2 deletion model of Rett Syndrome (1, 5, 17, 19, 20), we tested social interaction behaviors in each MeCP2 deletion line using the three-chamber sociability task (21). Control animals exhibited a significant preference for the chamber with the conspecific (C) over the empty chamber (E) (C: 50.7 ±2.4% E: 33.7 ± 3.7%; n = 18; p = 0.002; Wilcoxon matched pairs signed rank test; Fig. 5A). In contrast, the pan-interneuron Dlx5/6 mutants exhibited a reverse preference for the empty chamber (C: 26.1 ± 3.3% E: 47.6 ± 5.5%; n = 7; p = 0.015). We found no effect of MeCP2 deletion from PV interneurons on sociability (C: 49.8 ± 2.7% E: 26.8 ± 4.5%; n = 6; p = 0.03). However, SOM mutants showed no preference for either the conspecific or the empty chamber (C: 33.4 ± 2.6% E: 35.8 ± 4.0%; n = 13; p = 0.756) and the VIP mutants fully replicated the effects of the pan-interneuron deletion, showing a significant preference for the empty chamber over the conspecific (C: 27.2 ± 2.4% E: 43.9 ± 1.8%; n = 6; p = 0.031).

**Figure 5.**
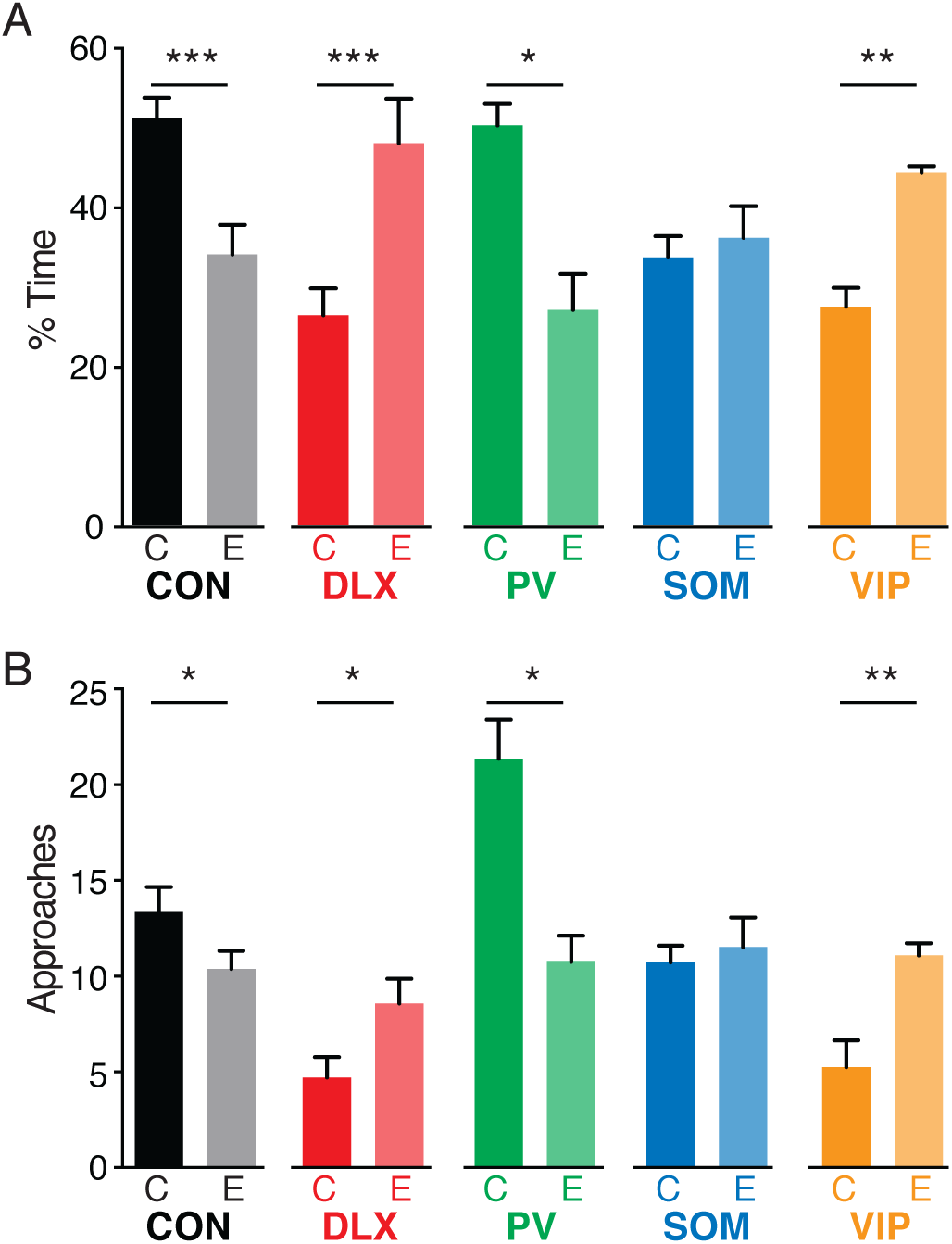
VIP and SOM interneurons contribute to impaired social behavior following MeCP2 deletion. A. Mean percentage of time spent with an unfamiliar conspecific (C) in a small holding cage versus an empty cage (E) for MeCP2^f/y^ controls (n = 14) and Dlx5/6 (n = 7), PV (n = 6), SOM (n = 13), and VIP (n = 6) mutants. B. Mean number of approaches made to within 5 cm of the conspecific or the empty holding cage for each group shown in A. *p < 0.05, ** p < 0.01, *** p < 0.001

In addition to altered overall social preferences, MeCP2 deletion from each interneuron class affected the number of approaches mice made towards conspecifics. Control animals made more approaches to the conspecific than to the empty holding cage (C: 14.3 ± 1.5 E: 10.4 ± 1.2; p = 0.030; Wilcoxon matched pairs signed rank test; Fig. 5B). In contrast, Dlx5/6 mutants made more approaches to the empty holding cage than the conspecific animal (C: 4.7 ± 1.1 E: 8.5 ± 1.3; p = 0.032). PV mutants showed a normal preference for approaching the conspecifics (C: 20.8 ± 2.1 E: 10.7 ± 1.4; p = 0.016), but made more approaches than did controls (C: 14.3 ± 1.5 E: 10.4 ± 1.2; p = 0.026). In contrast, SOM mutants made equal numbers of approaches to the conspecific and the empty holding cage (C: 10.6 ± 0.9 E: 11.4 ± 1.5; p = 0.55). Finally, the VIP mutants fully replicated the effects of pan-interneuron deletion, approaching the empty holding cage more than the conspecific (C: 5.2 ± 1.4 E: 11.0 ± 0.6; p = 0.023). Together, these data suggest that VIP interneurons, and potentially SOM interneurons, may contribute to the deficits in sociability behavior caused by global MeCP2 deletion.

## Discussion

Our results reveal a complex pattern of GABAeric contributions to behavioral phenotypes in the mouse model of Rett Syndrome and an unanticipated role for MeCP2 function in VIP interneurons. Based on previous characterizations of the MeCP2 deletion model (1, 2, 5, 17, 22), we examined interneuron contributions to three major categories of neural dysregulation: seizure and mortality, motor and anxiety behaviors, and social behaviors. We find evidence for overlapping contributions of all three major interneuron classes to seizure and motor dysfunction, but a unique role for MeCP2 deletion from VIP interneurons in social deficits.

Patients with Rett Syndrome exhibit respiratory impairments and seizure (1), and these phenotypes are replicated in mouse models following deletion of MeCP2 from all GABAergic cells (2, 5). We compared seizure and death incidence across interneuron cell type-specific MeCP2 loss of function mutations and a pan-interneuron loss of function. We found that MeCP2 deletion from each interneuron class increased the risk of seizure to some degree. In agreement with previous work (5), we found that MeCP2 loss of function in SOM interneurons conferred a late seizure phenotype beginning around postnatal day 200. However, conditional mutations of MeCP2 in the PV and VIP interneuron populations also caused modest increases in late seizure incidence. Seizure incidence was not associated with elevated mortality rates in the PV and VIP mutants, suggesting that the seizures were less severe than those in the SOM mutants. We likewise observed a low rate of late-onset seizure in the MeCP2^f/y^ control animals, indicating that the decrease in MeCP2 levels associated with the conditional allele (17) may be associated with epileptogenic consequences in addition to some mild behavioral phenotypes.

Distinct interneuron populations had different impacts on seizure incidence and overall mortality. Deletion of MeCP2 in the pan-interneuron Dlx5/6 mutants led to significant advances in seizure onset and mortality as compared to the SOM-specific mutants, suggesting a possible contribution from other interneuron populations such as PV cells. Because the PV promoter does not become active until ~2 weeks after birth, MeCP2 deletion from cells in the PV mutants likely did not occur until mid-adolescence (22). However, because the Dlx5/6 promoter turns on during early embryonic development in cells arising in both the medial and caudal ganglionic eminences (10, 11, 15), it is possible that early deletion of MeCP2 from PV interneurons in the Dlx5/6 animals promotes early seizure onset, as suggested for other developmental dysregulations of PV interneurons (15, 23, 24). Alternatively, deletion of MeCP2 from all three interneuron classes may have a supralinear impact on the early development of pathological activity patterns and respiratory impairments. A previous study did not observe seizure or early mortality phenotypes in the Dlx5/6 mutants, possibly due to methodological differences (2). MeCP2 knockout animals had significantly earlier seizure onset and mortality than the Dlx5/6 mutants, in agreement with previous work suggesting that loss of MeCP2 in excitatory neurons also contributes to seizure (25) and that respiratory impairments in the MeCP2 knockouts may increase early mortality (3, 4).

We further compared locomotor and anxiety phenotypes in each line and found that MeCP2 mutation in all three interneuron classes resulted in deficits in general motor function in the open field task, but only MeCP2 mutations in VIP interneurons led to deficits in marble burying, a task that is susceptible to altered anxiety and OCD-like behaviors as well as changes in fine motor function. None of the MeCP2 loss of function mutations resulted in anxiety-related phenotypes in the elevated plus maze, in agreement with previous work (2, 5). These data suggest that VIP interneuron dysfunction contributes to the motor deficits observed following broader deletion of MeCP2.

Abnormal or reduced social behavior is a hallmark of many autism spectrum disorder models, and has previously been shown in mice lacking MeCP2 in all GABAergic populations (2, 5). We found that pan-interneuron MeCP2 deletion mice exhibited a reversal of normal social preferences in the sociability assay, preferring an empty chamber to one containing a conspecific. Notably, MeCP2 deletion from VIP interneurons fully replicated this phenotype. In contrast, PV-specific deletion had no effect. Intriguingly, SOM-specific deletion led to loss of any social preference, suggesting a potential contribution of both VIP- and SOM-expressing cells to deficits in social behavior following pan-interneuron MeCP2 deletions. In agreement with results of a previous study using a partition test, which assays social preference between two conspecifics, we found that PV mutants showed elevated social interactions with novel conspecifics (5).

Our results comparing the PV and SOM interneuron contributions to seizure incidence and behavioral deficits are in agreement with some aspects of previous studies. However, we found very little impact of MeCP2 deletion in the PV-Cre mice. The PV-Cre line used here largely expresses in PV interneurons, along with some thalamocortical projection neurons (13, 26). In comparison, the PV-2A-Cre line used in some previous work (5) also expresses Cre in a subset of pyramidal neurons and additional thalamic nuclei (27). These differences may contribute to the more severe behavioral phenotypes and early mortality previously observed in the PV-2A-Cre line (5). Other work examining MeCP2 deletion in the PV-Cre line likewise found only mild behavioral phenotypes (22). However, as noted above, our findings from the PV-Cre line do not preclude a substantial contribution of embryonic MeCP2 deletion from PV interneurons to Rett Syndrome phenotypes.

Our findings suggest a partially overlapping set of contributions of different interneuron populations to seizure and behavioral deficits following deletion of MeCP2. In particular, we find an unanticipated and unique impact of MeCP2 deletion from the VIP interneurons. Despite being few in number (28), VIP interneurons are targets of multiple neuromodulatory systems and play critical roles in state-dependent regulation of local neural circuits (29-32), making them a potential point of neurodevelopmental vulnerability. The three interneuron classes examined here play key roles across many brain areas, including the cortex, hippocampus, and amygdala, and exhibit distinct cellular- and circuit-level properties. MeCP2 loss-of-function across multiple GABAergic interneuron classes may thus exert diverse influences on neural and behavioral deficits in Rett Syndrome.

## Acknowledgements

This work was funded by a Simons Foundation SFARI Pilot grant to J.A.C. The authors thank Victoria Hernandez and Brandon Wanke for assistance in performing behavioral assays and the lab of Dr. Jane Taylor for help with behavioral equipment.

## References

1. Chahrour M, Zoghbi HY (2007): The story of Rett syndrome: from clinic to neurobiology. Neuron. 56:422-437.

2. Chao HT, Chen H, Samaco RC, Xue M, Chahrour M, Yoo J, et al. (2010): Dysfunction in GABA signalling mediates autism-like stereotypies and Rett syndrome phenotypes. Nature. 468:263-269.

3. Chen RZ, Akbarian S, Tudor M, Jaenisch R (2001): Deficiency of methyl-CpG binding protein-2 in CNS neurons results in a Rett-like phenotype in mice. Nature genetics. 27:327-331.

4. Guy J, Hendrich B, Holmes M, Martin JE, Bird A (2001): A mouse Mecp2-null mutation causes neurological symptoms that mimic Rett syndrome. Nature genetics. 27:322-326.

5. Ito-Ishida A, Ure K, Chen H, Swann JW, Zoghbi HY (2015): Loss of MeCP2 in Parvalbumin-and Somatostatin-Expressing Neurons in Mice Leads to Distinct Rett Syndrome-like Phenotypes. Neuron. 88:651-658.

6. Shahbazian M, Young J, Yuva-Paylor L, Spencer C, Antalffy B, Noebels J, et al. (2002): Mice with truncated MeCP2 recapitulate many Rett syndrome features and display hyperacetylation of histone H3. Neuron. 35:243-254.

7. Amir RE, Van den Veyver IB, Wan M, Tran CQ, Francke U, Zoghbi HY (1999): Rett syndrome is caused by mutations in X-linked MECP2, encoding methyl-CpG-binding protein 2. Nature genetics. 23:185-188.

8. Hagberg B, Aicardi J, Dias K, Ramos O (1983): A progressive syndrome of autism, dementia, ataxia, and loss of purposeful hand use in girls: Rett’s syndrome: report of 35 cases. Annals of neurology. 14:471-479.

9. Ure K, Lu H, Wang W, Ito-Ishida A, Wu Z, He LJ, et al. (2016): Restoration of Mecp2 expression in GABAergic neurons is sufficient to rescue multiple disease features in a mouse model of Rett syndrome. eLife. 5.

10. Kepecs A, Fishell G (2014): Interneuron cell types are fit to function. Nature. 505:318-326.

11. Taniguchi H, He M, Wu P, Kim S, Paik R, Sugino K, et al. (2011): A resource of Cre driver lines for genetic targeting of GABAergic neurons in cerebral cortex. Neuron. 71:995-1013.

12. Monory K, Massa F, Egertova M, Eder M, Blaudzun H, Westenbroek R, et al. (2006): The endocannabinoid system controls key epileptogenic circuits in the hippocampus. Neuron. 51:455-466.

13. Hippenmeyer S, Vrieseling E, Sigrist M, Portmann T, Laengle C, Ladle DR, et al. (2005): A developmental switch in the response of DRG neurons to ETS transcription factor signaling. PLoS Biol. 3:e159.

14. Anderson SA, Eisenstat DD, Shi L, Rubenstein JL (1997): Interneuron migration from basal forebrain to neocortex: dependence on Dlx genes. Science. 278:474-476.

15. Wang Y, Dye CA, Sohal V, Long JE, Estrada RC, Roztocil T, et al. (2010): Dlx5 and Dlx6 regulate the development of parvalbumin-expressing cortical interneurons. J Neurosci. 30:5334-5345.

16. Zerucha T, Stuhmer T, Hatch G, Park BK, Long Q, Yu G, et al. (2000): A highly conserved enhancer in the Dlx5/Dlx6 intergenic region is the site of cross-regulatory interactions between Dlx genes in the embryonic forebrain. J Neurosci. 20:709-721.

17. Samaco RC, Fryer JD, Ren J, Fyffe S, Chao HT, Sun Y, et al. (2008): A partial loss of function allele of methyl-CpG-binding protein 2 predicts a human neurodevelopmental syndrome. Hum Mol Genet. 17:1718-1727.

18. Belzung C, Griebel G (2001): Measuring normal and pathological anxiety-like behaviour in mice: a review. Behav Brain Res. 125:141-149.

19. Kaufmann WE, Tierney E, Rohde CA, Suarez-Pedraza MC, Clarke MA, Salorio CF, et al. (2012): Social impairments in Rett syndrome: characteristics and relationship with clinical severity. J Intellect Disabil Res. 56:233-247.

20. Moretti P, Bouwknecht JA, Teague R, Paylor R, Zoghbi HY (2005): Abnormalities of social interactions and home-cage behavior in a mouse model of Rett syndrome. Hum Mol Genet. 14:205-220.

21. Nadler JJ, Moy SS, Dold G, Trang D, Simmons N, Perez A, et al. (2004): Automated apparatus for quantitation of social approach behaviors in mice. Genes Brain Behav. 3:303-314.

22. He LJ, Liu N, Cheng TL, Chen XJ, Li YD, Shu YS, et al. (2014): Conditional deletion of Mecp2 in parvalbumin-expressing GABAergic cells results in the absence of critical period plasticity. Nature communications. 5:5036.

23. Lau D, Vega-Saenz de Miera EC, Contreras D, Ozaita A, Harvey M, Chow A, et al. (2000): Impaired fast-spiking, suppressed cortical inhibition, and increased susceptibility to seizures in mice lacking Kv3.2 K+ channel proteins. J Neurosci. 20:9071-9085.

24. Rossignol E, Kruglikov I, van den Maagdenberg AM, Rudy B, Fishell G (2013): CaV 2.1 ablation in cortical interneurons selectively impairs fast-spiking basket cells and causes generalized seizures. Annals of neurology. 74:209-222.

25. Goffin D, Brodkin ES, Blendy JA, Siegel SJ, Zhou Z (2014): Cellular origins of auditory event-related potential deficits in Rett syndrome. Nat Neurosci. 17:804-806.

26. Cardin JA, Carlen M, Meletis K, Knoblich U, Zhang F, Deisseroth K, et al. (2009): Driving fast-spiking cells induces gamma rhythm and controls sensory responses. Nature. 459:663-667.

27. Madisen L, Zwingman TA, Sunkin SM, Oh SW, Zariwala HA, Gu H, et al. (2010): A robust and high-throughput Cre reporting and characterization system for the whole mouse brain. Nat Neurosci. 13:133-140.

28. Rudy B, Fishell G, Lee S, Hjerling-Leffler J (2011): Three groups of interneurons account for nearly 100% of neocortical GABAergic neurons. Developmental neurobiology. 71:45-61.

29. Fu Y, Tucciarone JM, Espinosa JS, Sheng N, Darcy DP, Nicoll RA, et al. (2014): A cortical circuit for gain control by behavioral state. Cell. 156:1139-1152.

30. Garcia-Junco-Clemente P, Ikrar T, Tring E, Xu X, Ringach DL, Trachtenberg JT (2017): An inhibitory pull-push circuit in frontal cortex. Nat Neurosci. 20:389-392.

31. Munoz W, Tremblay R, Levenstein D, Rudy B (2017): Layer-specific modulation of neocortical dendritic inhibition during active wakefulness. Science. 355:954-959.

32. Pi HJ, Hangya B, Kvitsiani D, Sanders JI, Huang ZJ, Kepecs A (2013): Cortical interneurons that specialize in disinhibitory control. Nature. 503:521-524.

